# Inferior Reinnervation of Reverse End-to-Side Nerve Transfer in a Delayed Nerve Repair Rat Model

**DOI:** 10.1101/2025.07.02.662849

**Authors:** Tak-Ho Chu, Amanda McConnachie, Oleksandra Kashyrina, Nicolas Lasaleta, Saud Alzahrani, Rajiv Midha

## Abstract

**Objective:** Reverse end-to-side (RETS) nerve transfer is a recent surgical technique to augment injured nerve function by supplying a dispensable donor motor nerve to the side of the distal injured nerve. Although clinical studies have suggested advantages of RETS transfer for upper extremity repairs, uncertainties remain regarding its underlying mechanism. Furthermore, our recent clinical studies using electrophysiological examinations revealed no contribution from the donor nerve. Given that most experimental studies were conducted on acutely injured nerves, our objective is to 1) reassess the effectiveness RETS nerve transfer in a rat model of chronic nerve injury and repair; 2) investigate the potential nerve-babysitting effect; and 3) investigate how availability of regenerating tracks, i.e. bands of Büngner, of recipient nerve affects donor nerve regeneration.

**Methods:** Obturator and femoral nerve were used as donor and recipient nerves, respectively. Electromyogram, retrograde labeling of regenerated motoneurons and neuromuscular junction (NMJ) formation were used to compare regenerative ability of donor nerve in acute and delayed RETS transfer where the femoral nerve in the latter group was injured by double ligations 8 weeks prior. Nerve-babysitting effect on injured nerve was investigated by 1) no intervention; 2) perineurial window creation; and 3) RETS transfer to femoral nerve in delayed repair model. The effects of availability of regeneration tracks were investigated by severing proximal femoral nerve, allowing complete denervation compared partial denervation in double ligations, followed by acute and delayed repairs.

**Results:** EMG and motoneuron quantification confirmed inferiority of donor nerve regeneration into recipient nerve in delayed RETS transfer compared to acute repair, yet donor axons reached target muscle and formed NMJs in both conditions. Same functional assessments revealed nerve baby-sitting effects did not significantly contribute to repair success but availability of regeneration tracks in the recipient nerve may influence the final outcomes.

**Conclusions:** Our study offered insights into the effectiveness of RETS nerve transfer in clinically relevant settings, underscoring the compounded impact of delayed intervention and native nerve regeneration which both negatively affect the efficacy of RETS nerve transfer.

## Introduction

Reverse end-to-side (RETS) nerve transfer was conceptualized two decades ago. The precursor model entailed the neurorrhaphy of transected tibial nerves to the side of transected peroneal nerves in rats and showed functional connection from the donor nerve to the end target ^1^. The same group further showed that following a proximal end-to-end repaired nerve, RETS transfer distal to the repair allows entry of donor axons to achieve functional recovery ^2^. These results sparked the enthusiasm towards RETS nerve transfer in experimental studies and eventually in clinical studies ^3,4^. Indeed, many case series demonstrated the efficacy of RETS transfer in upper extremity repairs ^5–7^. For example, as a recent randomized control trial provided evidence of the effectiveness of RETS in addition to ulnar nerve decompression and transposition alone in patients with chronic moderate-severe ulnar neuropathy at the elbow ^8^. However, unsatisfactory clinical findings also cast doubts on this repair technique ^9^. A multicentre non-randomized cohort study demonstrated significant improvement in key pinch strength in patients with RETS transfer, yet electrophysiological recording at the abductor digiti minimi muscle revealed no contribution from the donor anterior interosseous nerve three years after coaptation to a severely injured ulnar nerve ^10^.

Despite early adoption in nerve injury management, the exact mechanism of RETS nerve transfer is still not completely understood. Three possible roles of RETS transfer that can help with functional return are proposed: 1) axons from the donor nerve cross over to the injured recipient nerve and reinnervate the end target, which has been shown in rodent studies using axon histomorphometry and retrograde labeling ^1,2,4,11,12^; 2) the repair is primarily performed close to the muscle and reinnervating donor axons help preserve muscle health while allowing native axons from the proximal lesion to regenerate over a long distance, and reach the end target in a much later time, a similar concept borrowed from the “muscle-babysitting” technique ^13^; and 3) “nerve-babysitting” is believed to rejuvenate the injured nerve with trophic support from the donor nerve, causing an increase in the Schwann cell population in the recipient nerve at the coaptation site as demonstrated in a side-to-side bridge grafting model ^3,14^. In practice, the first two mechanisms would require donor axons to have successfully regenerated and navigated to the target muscle, and the last potential mechanism is yet to be confirmed.

Experimental rodent studies enable in depth analysis of axon regeneration under well controlled conditions. Our previous studies using peroneal-tibial nerve and obturator-femoral nerve in rodent RETS repair models demonstrated that creating a perineurial window maximized the opportunity for donor axons to navigate into the recipient nerve fascicle, supported by retrograde tracing and histological assessments ^12,15^. Other studies on RETS repair generally indicate comparable or superior efficacy of this transfer technique over other repair paradigms ^4,11,12,16,17^. However, most of the studies on RETS repair are performed at the same time as native nerve injury, the microenvironment within the distal recipient nerve is optimally permissive and inviting for regenerating axons. It is well-established that Schwann cells, upon the activation of the transcription factor cJun, adopt a repair phenotype after nerve injury, during which they clear myelin debris, proliferate, and secrete a plethora of growth factors, thereby creating a favourable environment for axonal growth ^18,19^. This phenotype declines with time, and Schwann cells gradually lose their capacity to support regeneration ^18,20,21^. It is possible that the beneficial effects of the RETS transfer technique are overestimated if only acute repair animal models are used. Therefore, a chronic experimental model of RETS may provide insights into the discrepancy between clinical outcome and lack of reinnervation from donor nerve in chronic neuropathy ^10^, when coupled with the use of viral labeling of donor axons, as well allow investigation into the potential mechanism of RETS repair under a chronic condition.

In the current study, we used a modified model that incorporated a two-month waiting period to create a chronically injured environment and traced donor axons and their reinnervation to target muscle. We hypothesized that a chronically injured recipient nerve is suboptimal for supporting the growth of regenerating axons from the donor nerve in delayed RETS nerve transfer. We also examined the role of nerve-babysitting on native nerve regeneration by comparing RETS transfer in a chronic condition with mechanical perturbation of the recipient nerve. Lastly, we compared the effects of partial and complete recipient nerve injury and determined how native axon regeneration affected donor axon regrowth.

## Materials and Methods

### Ethical Statement

All animal experiments were approved by the University of Calgary Animal Care Committee (AC20-0172). All applicable international, national, and institutional guidelines for the care and use of animals were followed.

### Surgical Procedures

Adult female Sprague-Dawley (SD) rats (225-250g, Charles River, Canada) were used. All animals were randomly assigned, and the surgeries were performed under deep anesthesia maintained with 2% isoflurane using aseptic technique. Preoperative analgesics included buprenorphine (0.05 mg/kg, s.c.) and meloxicam (1 mg/kg, s.c.); and post-operative analgesic meloxicam (1 mg/kg, s.c.) were given for 2 days after surgeries. To compare acute and delayed repair, acute RETS nerve transfers (n=8) were performed as previously described ^15^. Briefly, proximal femoral nerve (FN) proper was crushed and double ligated with 10-0 sutures, obturator nerve (ON) was then mobilized and inserted into the femoral nerve motor branch (MFN) through a perineurial opening and secured using 10-0 sutures. For delayed repair (n=8), proximal FN was double ligated and waited for two months before RETS transfer. Electromyography (EMG) were performed at five weeks after RETS transfer by stimulating at FN and recording at the rectus femoris muscle with a single supramaximal stimulation at 2.5mA with 50 µs pulse width. FN was then cut to evaluate contribution from ON only. Onset latency and peak amplitude were acquired and averaged from three stimulations at different locations at the muscle. Additional EMG data (n=10 in acute and n=9 in delayed repair) were added from a pilot study which had same surgical procedures but omitted from other analyses due to technical error in labeling (injection of tracers using Hamilton syringe causing over-estimation of regenerated motor neurons). After electrophysiological recordings, MFN was cut distally before muscle entry, the proximal cut end of MFN and proximal cut end of FN were soaked in retrograde tracers micro-ruby and micro-emerald, respectively, using gel foam in silicone tubes for one hour to allow tracer uptake. The animals were kept for one week before tissue harvest.

To determine the effect of “nerve babysitting” and lesioning at the coaptation site on native FN regeneration, FN was ligated for two months before the animals were random assigned into three groups: 1) ON cut (n=7); 2) ON cut and perineurial window created (n=9); and 3) ON RETS transfer (n=8). EMG were performed as described above but only FN was stimulated and ON was cut from coaptation in animals with ON RETS transfer to determine the contribution from FN alone. Retrograde labeling was performed as above except without cutting FN and only micro-ruby was used at the MFN.

To determine the effect of acute and chronic complete denervation at the recipient nerve, animals were randomly assigned into two groups: 1) FN cut and immediate RETS transfer (n=6); 2) FN cut and delayed RETS transfer after two months (n=6). Distal cut end of FN was sutured onto adjacent abdominal wall muscle with 9-0 suture. EMG were performed five weeks after transfer as described above but only ON was stimulated. Retrograde labeling was performed as above except micro-ruby was used at the MFN.

To determine donor nerve contribution to neuromuscular junction (NMJ) formation in acute and delayed nerve repair, two groups of animals receiving acute (n=7) and delayed (n=7) repairs were intraneurally injected with 0.5ul of pAAV-CAG-GFP (PHPeB, Addgene, USA) into the obturator nerve prior to RETS transfer using a 34-gauge Hamilton syringe. Animals were kept for two weeks (n=2 in each group) and six weeks (n=5 in each group) to visualize green fluorescent protein (GFP) positive axons in recipient nerve and target muscle respectively. No EMG or labeling was performed.

### Perfusion, tissue clearing, and imaging

At the end of the survival period, animals were euthanized with overdose sodium pentobarbital (Euthanyl) and perfused intracardially with 4% paraformaldehyde. Spinal cords were harvested, cleared using gradient of tetrahydrofuran and ethyl cinnamate and the whole lumbar spinal cord were imaged with lightsheet microscopy (UltraMicroscope II, Miltenyi, Germany) as previously described ^22^. Retrograde labelled motoneurons were counted using automated algorithm in imaging software Imaris (v10, Oxford Instruments, UK).

### Muscle and nerve sectioning, staining, imaging, and neuromuscular junction quantification

For animals in NMJ quantification groups, rectus femoris muscles were harvested six weeks after nerve transfer and processed for frozen sectioning and immunostaining as previously described ^22^. Briefly, nerves and muscles were embedded in OCT and cut in longitudinal sections at a thickness of 20 μm on a cryostat. For immunostaining, a mixture of anti-neurofilament (SMI312, Biolegend, USA) and anti-synaptic vesicles (SV2, developmental studies hybridoma bank, USA) was used to label regenerating axons, and Alexa-555 conjugated α-bungarotoxin (Invitrogen, USA) was used to label NMJs. Sections were mounted with Fluoromount mounting medium and imaged with a slide scanner (VS110, Olympus, Japan). Every tenth section of the muscle block was analyzed and the number of *en-face* un-occupied NMJs, NMJs with NF + SV2 labeling and NMJs with GFP labeling were quantified manually on QuPath ^23^ by a blinded observer.

### Statistical Analysis

Data were analyzed with Prism 10 software (GraphPad, USA) using Student’s *t*-test, one-way or two-way ANOVA and post hoc Tukey’s multiple comparison test, where appropriate, to compare results between groups. Statistical significance was accepted at *p* < 0.05, with results present as the mean ± SEM.

## Results

### Effects of acute vs delayed RETS nerve transfer

We first determined the effects of acute and delayed RETS nerve transfer on axon regeneration and muscle reinnervation using EMG and retrograde labeling of regenerated motor neurons (Fig 1A). While recording at the rectus femoris muscle, stimulating the FN resulted in a significantly shorter onset latency and a higher peak amplitude in the delayed repair, indicating significant regrowth of native FN axons over time (Fig 1B). When stimulating at the ON after FN transection, the peak amplitude was significantly higher in the acute repair, indicating greater input from donor axons, whereas onset latency remained similar (Fig 1C). Retrograde labeling at the proximal FN showed a significantly higher number of motor neurons breaching the ligations in the delayed compared to the acute repair (Fig 1D, micro-emerald), whereas labeling at the MFN (with the FN transected, Fig 1D, micro-ruby) showed a significantly higher number of motor neurons in the acute compared to the delayed repair, suggesting more ON axons regenerated into the recipient nerve in the acute repair.

**Fig 1.**
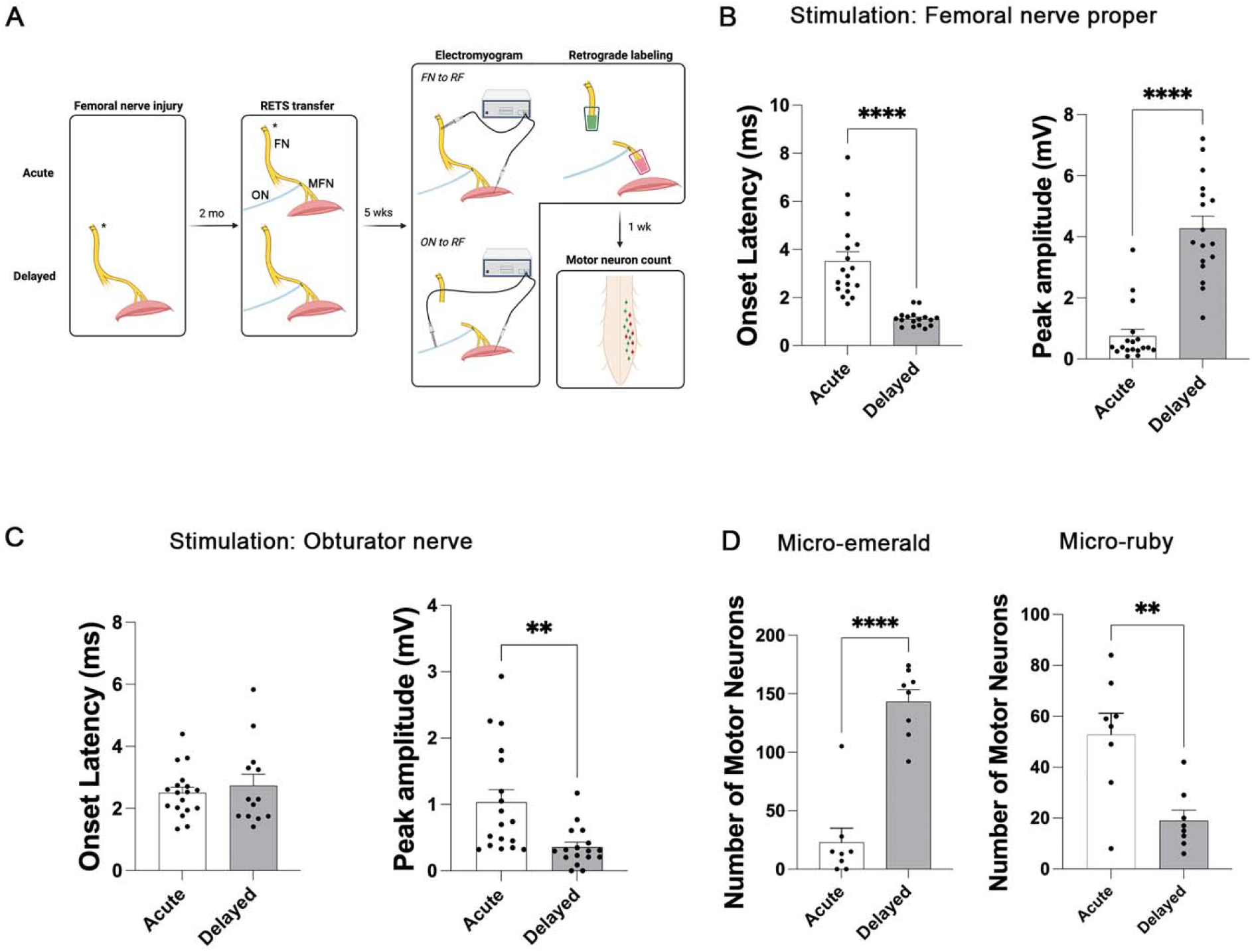
Superior regeneration in acute RETS repair compared to delayed repair. (A) Schematic diagram showing experimental procedures. Femoral nerve (FN) injury was created by double crush and ligations with 10-0 sutures (asterisk), obturator nerve (ON, blue) was coapted to femoral nerve motor branch (MFN) immediately or delayed for 2 months, sensory branches of FN are shaded. Electromyogram (EMG) and retrograde labeling were performed 5 weeks after coaptation and spinal cords were harvested 1 week after. Retrograde labeled motor neurons were then counted. (B) Stimulation at FN proper and recording at rectus femoris (RF) showed significantly longer onset latency and lower peak amplitude in acute repair compared to delayed or delayed ON repairs. (C) Stimulation at ON and recording at RF showed similar onset latency but significantly higher peak amplitude in acute repair compared to delayed ON repairs. Data from B and C are combined from current study and a previous pilot cohort. (D) Quantification of retrograde labeled motor neurons showed significantly more FN motor neurons (micro-emerald positive) breached the ligations in delayed repairs but more ON motor neurons (micro-ruby positive) regenerated into the recipient nerve in acute as compared to delayed repair. **, p<0.01; ***, p<0.001; ****. p<0.0001. Student’s t-test.

### Nerve and muscle reinnervation in acute vs delayed RETS nerve transfer

To examine whether donor axons reinnervate the recipient nerve and target muscle, we injected an adeno-associated virus (AAV, PHP.eB-GFP) that predominantly transduces neurons into the ON to distinguish donor axons from native FN axons (Fig 2A). Motor neuron counts showed high efficiency with no statistical difference in the number of transduced motor neurons between groups, averaging 88 and 70 in the acute and delayed repair groups, respectively, out of an average of 98 ON motor neurons as previously described ^15^. Immunostaining of the coapted nerve two weeks after RETS transfer showed an abundance of neurofilaments in the delayed repair, whereas ON axons expressing GFP were found in both the acute and delayed repair groups with bidirectional growth (Fig 2B, white arrows), similar to our previous findings ^15^. Additional animals survived six weeks after RETS transfer, their muscles were harvested, immunostained with a combination of neurofilament and synaptic vesicle 2, and subsequently labeled with *α*-bungarotoxin, which recognizes NMJs. Results showed reinnervation by GFP positive donor axons in both acute and delayed repair conditions (Fig 2C, arrows). Quantification of different innervation states of NMJs revealed a similar number of denervated and native axon innervated NMJs in both conditions (Fig 2D). However, the number of NMJs reinnervated by donor axons was significantly higher in the acute repair group compared to the delayed repair group.

**Fig 2.**
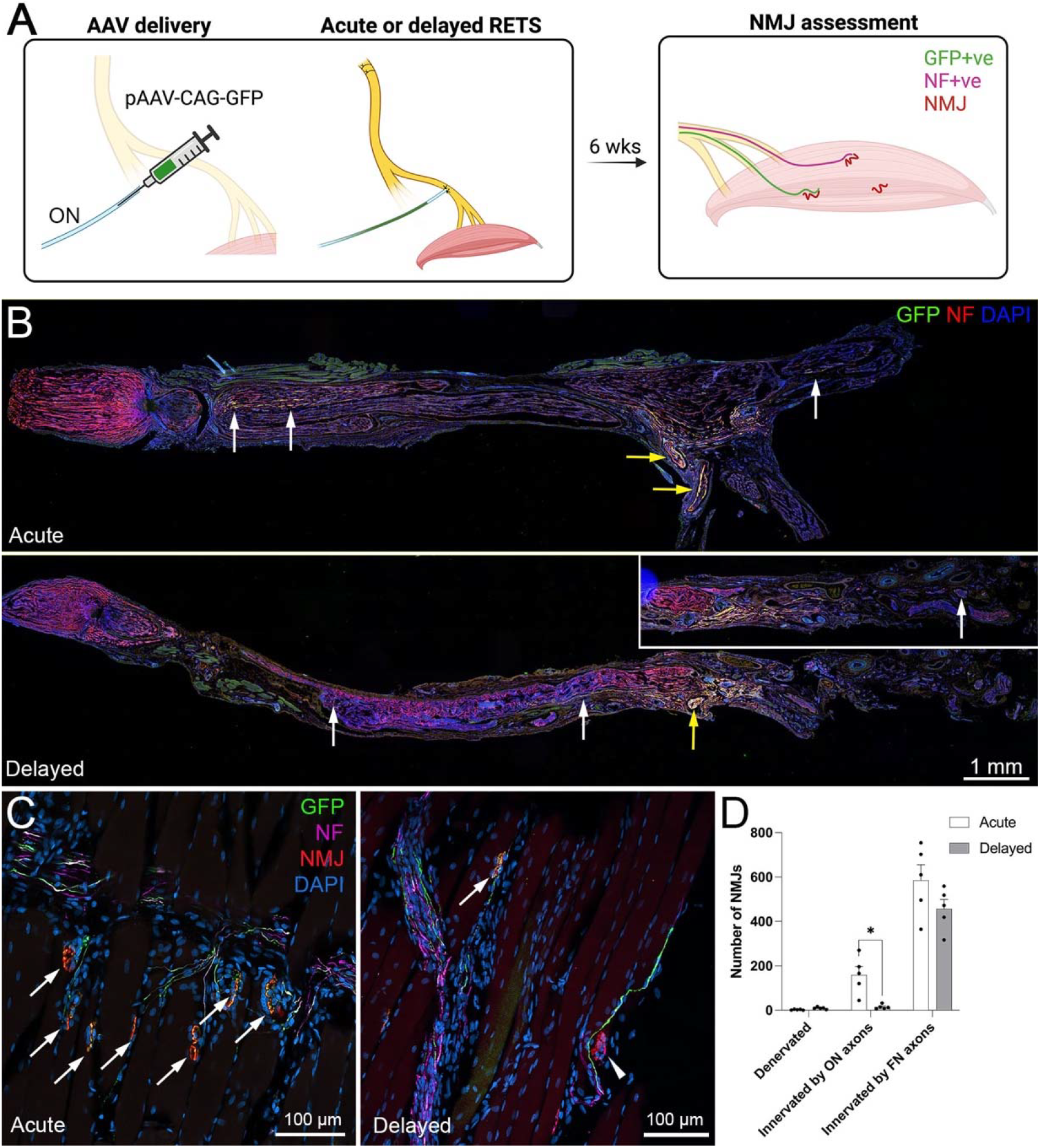
Neuromuscular junctions (NMJs) were reinnervated by donor axons in acute and delayed nerve transfer. (A) Schematic diagram showing injection of ON with neuron specific adenovirus-associated virus before coaptation to MFN in acute or delayed RETS transfer. Nerve samples were examined at 2 weeks (not shown) and muscle samples were examined at 6 weeks for NMJ reinnervation. (B) Slide scanner images of coapted nerve complexes showing GFP positive axons (white arrows) from ON (yellow arrows) regenerating in both directions (proximal on left) after entering recipient nerve. More neurofilament positive axons were found in delayed repair. (C) Representative images showing NMJs reinnervation by GFP donor nerve axons in acute and delayed repair animals (arrows). Arrowhead points to a denervated NMJ. (D) Quantification of denervated, innervated by GFP positive, and neurofilament positive NMJs revealed significantly more NMJs innervated by donor axons in acute compared to delayed repair. *, p<0.05. Two-way ANOVA.

### Effects of nerve babysitting

One potential mechanism proposed for RETS nerve transfer is the rejuvenation of the recipient nerve via Schwann cells or growth factors from the donor nerve ^3^, which then promotes native axon regeneration. We examined this hypothesis by comparing delayed RETS transfer with sham procedures without MFN lesioning (ON cut only) or with perineurial window creation at the MFN (ON cut + perineurial window (PW)) to examine the effect of perturbation on the recipient FN (Fig 3A). All animals received FN double ligations two months prior to intervention when the ON was cut. Animals were kept for five weeks, and EMG and retrograde labeling were subsequently performed. While recording at the rectus femoris muscle, stimulating the FN showed no difference among all groups in onset latency and peak amplitude (Fig 3B). Retrograde labeling at the MFN also showed no difference across all groups, suggesting that neither the presence of a donor nerve or manipulation of the recipient nerve with a perineurial window influenced native axon regeneration (Fig 3B).

**Fig 3.**
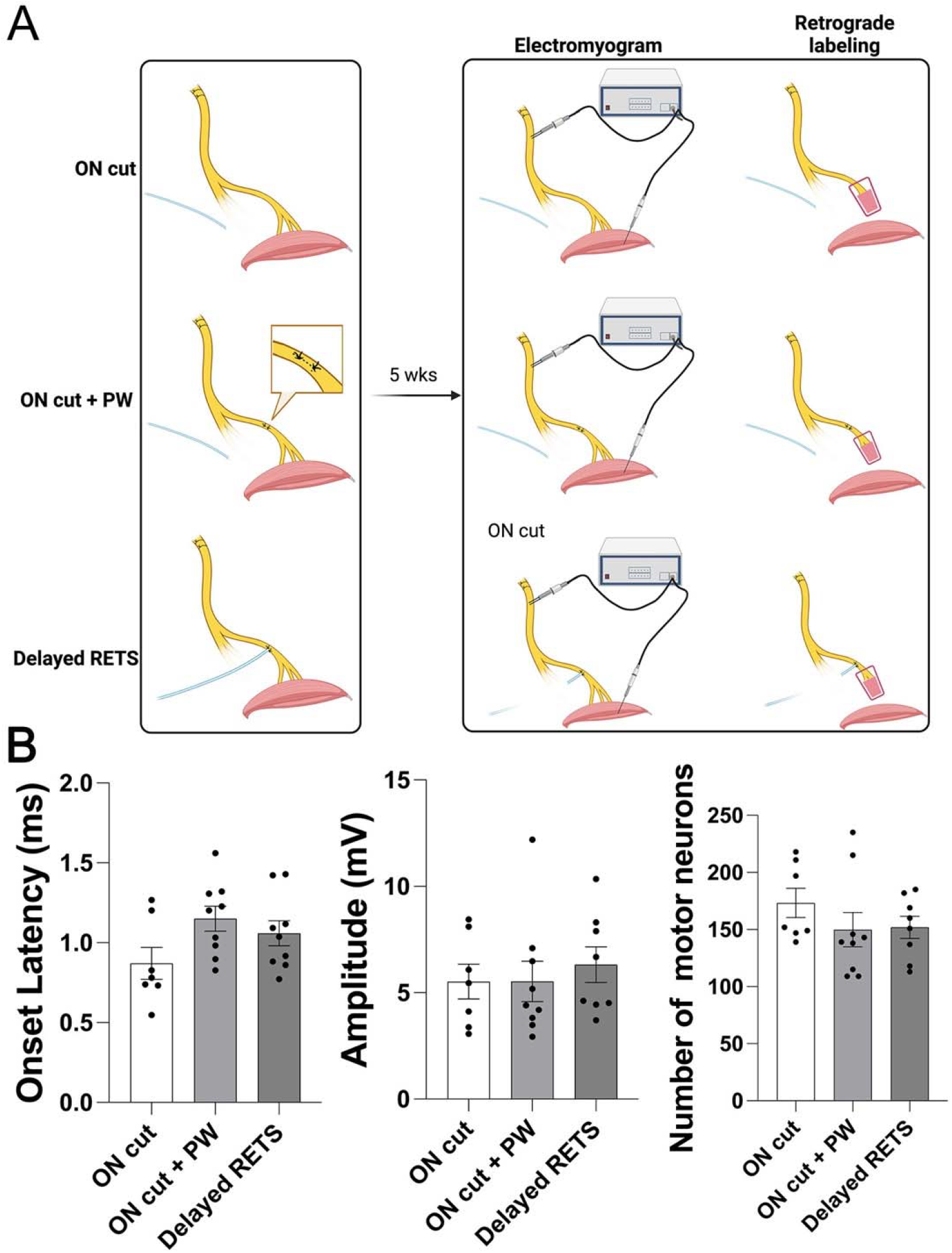
RETS nerve transfer did not contribute to nerve babysitting. (A) schematic diagram showing experimental procedures. Femoral nerve was double ligated in all groups two months before receiving additional intervention, one group received ON cut, one group received ON cut with perineurial window (PW) opening and closed with sutures, one group received RETS nerve transfer. EMG and retrograde labeling were performed 5 weeks after interventions. (B) Onset latency, peak amplitude and motor neuron counts were comparable between all groups using one-way ANOVA.

### Effects of acute repair vs delayed repair in a complete denervated nerve

Since double ligations of the proximal FN did not prevent native axons from regenerating into the distal nerve, which might compete with the reinnervating donor axons for available Schwann cell tracks, we therefore transected the nerve to enable complete denervation before performing acute or delayed RETS transfer (Fig 4A). EMG and motor neuron counts (Fig 4B) revealed similar recovery kinetics from the donor nerve, with no significant difference in onset latency, peak amplitude, or number of retrogradely labeled motor neurons between the acute and delayed repair groups.

**Fig 4.**
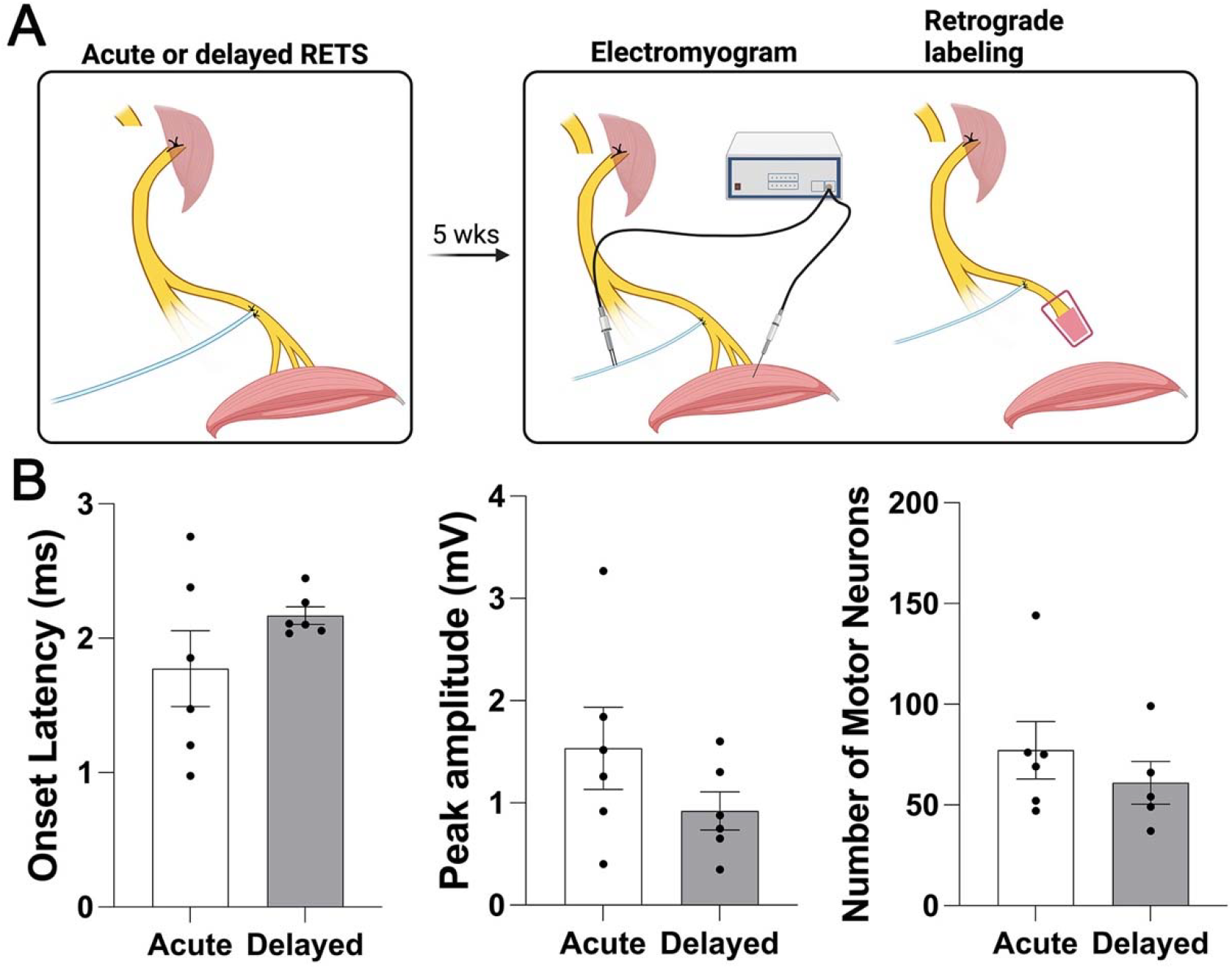
Similar regeneration profile when recipient nerve is completely denervated. (A) schematic diagram showing experimental procedures. Femoral nerve was transected before acute or delayed RETS transfer. EMG and retrograde labeling were performed 5 weeks after interventions. (B) Onset latency, peak amplitude and motor neuron counts were comparable between acute and delayed repair groups using Student’s t-test.

## Discussion

To the best of our knowledge, all experimental RETS transfer studies in the literature were conducted in an acute setting where a donor nerve was transferred to a freshly injured recipient nerve. This scenario is clinically rare and likely skewed the results due to the presence of proliferative repair Schwann cells in the recipient nerve days after the injury. Our chronic injury model provides insights into the mechanism of RETS and how the transfer technique can be further refined in pre-clinical models. Our findings have clinical implications for patient selection, such as for the anterior interosseous nerve (AIN)-to-ulnar motor RETS nerve transfer procedure, which is now common in clinical settings ^24^.

A comparison between acute and delayed RETS repair (Fig 1 and Table 1) showed a clear advantage of acute repair, as evidenced by the significantly higher peak amplitude (Fig 1C) and a greater number of micro-ruby positive motor neurons (53 ± 8), indicating donor nerve motor neurons that had regenerated into the distal recipient FN (Fig 1D). This corresponds to slightly more than half (54.1%) of the total ON motor neuron pool at the level used for coaptation (averaging 98 motor neurons from previous assessments ^15^). However, in chronic conditions, only one-fifth (19.4%) of ON motor neurons (19 ± 4) were able to regenerate into the distal recipient nerve. EMG data also showed that the conductivity of the donor nerve in delayed repair reached only 40% of the peak amplitude achieved by that in acute repair (0.41 *vs* 1.04 mV, Fig 1C and Table 1). In contrast, native FN axons were able to breach the double ligations and recover to 33% of baseline amplitude (4.28 *vs* 13.1 mV from the contralateral side) by 13 weeks in delayed repair (Fig 1B). This aligns with 41.1% of micro-emerald-labeled motor neurons (143 ± 10) out of the total FN motor neuron pool (averaging 348 motor neurons from previous assessments ^15^). In the acute repair condition, native FN axons were only able to recover to 6% of baseline amplitude (0.75 *vs* 13.1 mV) and 6.6% of FN motor neuron pool (23 ± 12) had regenerated into the distal motor branch by five weeks at the time of labeling. The data suggest robust native FN axon regeneration over time despite physical disruption to the nerve structure (double ligations). It is evident that some axons grew outside of the perineurium at the ligation sites to achieve the breach (Fig 2B).

The “nerve-babysitting” effect was not observed when a donor nerve was coapted to the recipient nerve compared to no coaptation or only perturbation at the distal nerve by creating a perineurial window (Fig 3B). In fact, the transected proximal end of the donor nerve can provide only a minimal number of repair Schwann cells, since Wallerian degeneration and Schwann cell activation are only initiated in the nerve segment distal to the lesion ^18^. It is likely that physical perturbation (perineurial window creation) at the coaptation site generated a portion of repair Schwann cells locally, which can stimulate native nerve outgrowth. However, the beneficial effect is perhaps negated by the damage of already regenerated axons (Fig 3B). Therefore, it remains to be determined whether preconditioning by crushing the donor nerve proximally would induce a meaningful amount of repair Schwann cells to migrate into the recipient nerve and further promote recipient nerve axon regeneration.

Since half of the recipient FN motor neurons regenerated into the distal nerve, the double crush and ligations at the proximal FN effectively functioned as a nerve injury in continuity model. The condition is not ideal since such level of reinnervation from native nerve would not necessitate nerve transfer. To investigate how the innervation status of recipient nerve affects donor nerve regeneration in chronic setting, we therefore transected the proximal FN to create a completely denervated recipient nerve. We observed a trend, yet statistically insignificant, of reduced peak amplitude and labeled motor neuron count in delayed repair after transection (Fig 4). When comparing partial FN injury with crush and double ligations versus complete denervated FN injury with transection, EMG metrics, though not statistically significant, were trending lower in onset latency and higher in peak amplitude in the complete denervation groups, regardless of acute or delayed repair (Supp Fig 1A & B). Similarly, a trend of more ON motor neurons regenerated into the distal recipient nerve in both acute and delayed repair following transection (78.5% and 62.2% of the total ON motor neuron pool, respectively) compared to partial FN injury (Supp Fig 1C). In fact, significantly more motor neurons regenerated into the transected recipient FN compared to the partially injured FN under delayed conditions. This strongly suggests that besides the chronic status of the recipient nerve, the availability of regeneration tracks is also an important confounding factor. This further consolidates our previous study ^12^ which demonstrated the negative effect of donor axons on native regeneration, as donor axons were able to pre-emptively regenerate into and occupy the regeneration tracks in the more distal segment of the recipient nerve. Recent rodent studies examining axonal growth into an intact, naïve nerve using a sensory nerve bridge approach ^22^ or an upper arm model of RETS ^25^ confirmed the inability of donor motor axons to reach the recipient nerve when no regeneration tracks were available. We speculate that the same mechanism applies to delayed repair in which the regeneration tracks were already occupied by the native axons at two months after injury, thereby hampering successful regeneration of donor axons at the time of coaptation.

To further examine the functional connection between the donor nerve and muscle reinnervation, we labeled the donor nerve motor neuron pool with neurotropic AAV-PHP.eB, an AAV9 variant that specifically transduces neurons of the central and peripheral nervous systems ^26^. Previous reports showed that intravenous injection of AAV-PHP.eB efficiently labeled spinal motor neurons ^27^. Our results demonstrated that injection of AAV-PHP.eB expressing a GFP transgene into the ON resulted in strong GFP expression in spinal motor neurons, dorsal root ganglion neurons (data not shown), and their projections (Fig 2B & C). The technique enabled unambiguous identification of donor axon reinnervation in motor endplates. Confocal microscopy confirmed the presence of NMJs reinnervated by donor axons in both conditions (Fig 2C). Similar to previous studies ^28^, the number of denervated NMJs significantly decreased over time and was minimal at either 6 weeks (acute) or 14 weeks (delayed) after double ligations (Fig 2D). The significantly higher number of NMJs reinnervated by donor axons in the acute group aligns with a higher peak amplitude in CMAP recording when the obturator nerve was stimulated (Fig 1C). However, the number of NMJs innervated by native axons in acute group is unexpectedly high, although it is possible that some may represent donor axons not successfully transduced with the injected AAVs, the more likely possibility is that some degenerated axonal fragments from the original innervation were not completely removed and mistakenly over-estimated as reinnervation by native axons.

Overall, our EMG, retrograde labeling and histological assessments demonstrated that while only a small portion of donor axons successfully regenerated into the coapted nerve, they were able to reinnervate the target muscles, even under chronic conditions. These findings align with a recently published randomized controlled trial that demonstrated the effectiveness of AIN-to-ulnar motor nerve RETS transfer in chronic cases, even after more than six months post-symptom onset ^8^. Patients who underwent nerve transfer exhibited superior outcomes, including improved grip strength, larger CMAPs, better Disabilities of the Arm, Shoulder, and Hand (DASH) scores, and greater reductions in muscle atrophy compared to those who received conventional surgery ^8^. Our rodent study was limited by the use of an eight-week period to establish chronic conditions. Previous research has shown that axon regeneration is significantly impaired if denervation exceeds four weeks in rats ^29^. Additionally, marker genes for denervated Schwann cells, such as p75 and cJun, peak one week after nerve injury in mice and are down-regulated after six and three weeks, respectively ^30^. Recent studies in mice also revealed that repair Schwann cells transition to senescent Schwann cells six weeks after nerve transection. These senescent cells exhibit a senescence-associated secretory phenotype, characterized by pro-inflammatory angiogenic and extracellular matrix degrading factors that ultimately inhibit axonal growth ^31^. Other rodent studies have indicated that while axon regeneration is still possible after three to six month of distal nerve denervation following end-to-end repair with a heathy proximal nerve, functional recovery is entirely absent ^32^. Therefore, it is plausible that our results for delayed repair would have been less favourable if a longer wait time had been used to establish chronic conditions. Yet, it also highlights the temporal differences in the degenerative process and recovery kinetics between rodents and humans ^33^, with the latter exhibiting a wider pro-regenerative window of up to 100 days post-injury, during which cJun, p75, and Sox10 begin to decline in human nerves ^34^.

## Conclusions

Our results demonstrated that a small portion of donor nerve axons regenerated and reinnervated target muscles after RETS nerve transfers especially in chronic conditions in which outcomes were inferior to acute repair. Our study underscores the compounded impact of delayed intervention and occupancy of regeneration tracks by native axons, both of which negatively affect the efficacy of RETS nerve transfer.

## Acknowledgments

The study is partially supported by the Plastic Surgery Foundation Combined Pilot Research Grant awarded to THC and RM. We acknowledge the Hotchkiss Brain Institute Advanced Microscopy Platform and the Cumming School of Medicine for support and use of their confocal, slide scanner, and light-sheet microscope. Schematic figures were created with Biorender.com.

## References

1. Isaacs J, Allen D, Chen LE, Nunley J. Reverse End-to-Side Neurotization. J Reconstr Microsurg. 2005;21(01):43–48. doi:10.1055/s-2005-862780

2. Isaacs J, Cheatham S, Gagnon E, Razavi A, McDowell C. Reverse End-to-Side Neurotization in a Regenerating Nerve. J Reconstr Microsurg. 2008;24(07):489–496. doi:10.1055/s-0028-1088230

3. Isaacs J. Reverse End-to-Side (Supercharging) Nerve Transfer: Conceptualization, Validation, and Translation. Hand. 2022;17(6):1017–1023. doi:10.1177/1558944720988076

4. Kale SS, Glaus SW, Yee A, et al. Reverse End-to-Side Nerve Transfer: From Animal Model to Clinical Use. J Hand Surg Am. 2011;36(10):1631–1639.e2. doi:10.1016/j.jhsa.2011.06.029

5. Davidge KM, Yee A, Moore AM, Mackinnon SE. The Supercharge End-to-Side Anterior Interosseous–to–Ulnar Motor Nerve Transfer for Restoring Intrinsic Function: Clinical Experience. Plast Reconstr Surg. 2015;136(3). https://journals.lww.com/plasreconsurg/fulltext/2015/09000/the_supercharge_end_to_side_anterior.18.aspx

6. Baltzer H, Woo A, Oh C, Moran SL. Comparison of Ulnar Intrinsic Function following Supercharge End-to-Side Anterior Interosseous-to-Ulnar Motor Nerve Transfer: A Matched Cohort Study of Proximal Ulnar Nerve Injury Patients. Plast Reconstr Surg. 2016;138(6):1264–1272. doi:10.1097/PRS.0000000000002747

7. Pathiyil RK, Alzahrani S, Midha R. Reverse End-to-Side Transfer to Ulnar Motor Nerve: Evidence From Preclinical and Clinical Studies. Neurosurgery. 2023;92(4). https://journals.lww.com/neurosurgery/fulltext/2023/04000/reverse_end_to_side_transfer_to_ulnar_motor_nerve_.4.aspx

8. Xie Q, Shao X, Song X, et al. Ulnar nerve decompression and transposition with versus without supercharged end-to-side motor nerve transfer for advanced cubital tunnel syndrome: a randomized comparison study. J Neurosurg. 2022;136(3):845–855. doi:10.3171/2021.2.JNS203508

9. Thorkildsen RD, Kleggetveit IP, Thu F, Madsen LM, Bolstad BJ, Røkkum M. Supercharging of the ulnar nerve: clinical and neurophysiological assessment at 2 years for nine proximal injuries. Journal of Hand Surgery (European Volume). 2024;49(9):1139–1146. doi:10.1177/17531934231226174

10. Curran MWT, Olson JL, Morhart MJ, et al. Reverse End-to-Side Nerve Transfer for Severe Ulnar Nerve Injury: A Western Canadian Multicentre Prospective Nonrandomized Cohort Study. Neurosurgery. 2022;91(6). https://journals.lww.com/neurosurgery/Fulltext/2022/12000/Reverse_End_to_Side_Nerve_Transfer_for_Severe.5.aspx

11. Fujiwara T, Matsuda K, Kubo T, et al. Axonal supercharging technique using reverse end-to-side neurorrhaphy in peripheral nerve repair: An experimental study in the rat model. J Neurosurg. 2007;107(4):821–829. doi:10.3171/JNS-07/10/0821

12. Nadi M, Ramachandran S, Islam A, Forden J, Guo GF, Midha R. Testing the effectiveness and the contribution of experimental supercharge (reversed) end-to-side nerve transfer. J Neurosurg. Published online 2018. doi:10.3171/2017.12.JNS171570.

13. Terzis JK, Tzafetta K. The “Babysitter” Procedure: Minihypoglossal to Facial Nerve Transfer and Cross-Facial Nerve Grafting. Plast Reconstr Surg. 2009;123(3). https://journals.lww.com/plasreconsurg/fulltext/2009/03000/the_babysitterprocedure_minihypoglossal_to.12.aspx

14. Isaacs J, Feger MA, Mallu S, et al. Side-to-side supercharging nerve allograft enhances neurotrophic potential. Muscle Nerve. 2020;61(2):243–252. doi:10.1002/mus.26753

15. Chu TH, Alzahrani S, McConnachie A, et al. Perineurial Window is Critical for Experimental Reverse End-to-Side Nerve Transfer. Neurosurgery. 2023;93(4):952–960. doi:10.1227/neu.0000000000002481

16. von Guionneau N, Sarhane KA, Brandacher G, Hettiaratchy S, Belzberg AJ, Tuffaha S. Mechanisms and outcomes of the supercharged end-to-side nerve transfer: a review of preclinical and clinical studies. J Neurosurg. 2021;134(5):1590–1598. doi:10.3171/2020.3.JNS191429

17. Farber SJ, Glaus SW, Moore AM, Hunter DA, Mackinnon SE, Johnson PJ. Supercharge nerve transfer to enhance motor recovery: A laboratory study. Journal of Hand Surgery. 2013;38(3):466–477. doi:10.1016/j.jhsa.2012.12.020

18. Jessen KR, Mirsky R. The repair Schwann cell and its function in regenerating nerves. Journal of Physiology. 2016;594(13):3521–3531. doi:10.1113/JP270874

19. Arthur-Farraj PJ, Latouche M, Wilton DK, et al. c-Jun Reprograms Schwann Cells of Injured Nerves to Generate a Repair Cell Essential for Regeneration. Neuron. 2012;75(4):633–647. doi:10.1016/j.neuron.2012.06.021

20. Höke A, Gordon T, Zochodne DW, Sulaiman OAR. A Decline in Glial Cell-Line-Derived Neurotrophic Factor Expression Is Associated with Impaired Regeneration after Long-Term Schwann Cell Denervation. Exp Neurol. 2002;173(1):77–85. doi:10.1006/exnr.2001.7826

21. Fu SY, Gordon T. Contributing factors to poor functional recovery after delayed nerve repair: prolonged denervation. The Journal of Neuroscience. 1995;15(5):3886LP–3895. doi:10.1523/JNEUROSCI.15-05-03886.1995

22. Alzahrani S, Brito da Silva H, Chu T ho, et al. Successful retrograde regeneration using a sensory branch for motor nerve transfer. J Neurosurg. Published online July 1, 2022:1–10. doi:10.3171/2022.6.JNS22734

23. Bankhead P, Loughrey MB, Fernández JA, et al. QuPath: Open source software for digital pathology image analysis. Sci Rep. 2017;7(1):16878. doi:10.1038/s41598-017-17204-5

24. Chambers SB, Wu KY, Ross DC, Gillis JA. Anterior Interosseus to Ulnar Motor Nerve Transfers: A Canadian Perspective. HAND. 2023;19(7):1075–1079. doi:10.1177/15589447231174482

25. Harnoncourt L, Schmoll M, Festin C, et al. Axonal regeneration and innervation ratio following supercharged end-to-side nerve transfer. 2025;(February):1–11. doi:10.3389/fcell.2025.1513321

26. Chan KY, Jang MJ, Yoo BB, et al. Engineered AAVs for efficient noninvasive gene delivery to the central and peripheral nervous systems. Nat Neurosci. 2017;20(8):1172–1179. doi:10.1038/nn.4593

27. Dayton RD, Grames MS, Klein RL. More expansive gene transfer to the rat CNS: AAV PHP.EB vector dose–response and comparison to AAV PHP.B. Gene Ther. 2018;25(5):392–400. doi:10.1038/s41434-018-0028-5

28. Vannucci B, Santosa KB, Keane AM, et al. What is Normal? Neuromuscular junction reinnervation after nerve injury. Muscle Nerve. 2019;60(5):604–612. doi:10.1002/mus.26654

29. Sulaiman OAR, Gordon T. Effects of short- and long-term Schwann cell denervation on peripheral nerve regeneration, myelination, and size. Glia. 2000;32(3):234–246. doi:10.1002/1098-1136(200012)32:3<234::AID-GLIA40>3.0.CO;2-3

30. Wagstaff LJ, Gomez-Sanchez JA, Fazal S V, et al. Failures of nerve regeneration caused by aging or chronic denervation are rescued by restoring Schwann cell c-Jun. Chao MV, Bronner ME, Hoke A, eds. Elife. 2021;10:e62232. doi:10.7554/eLife.62232

31. Fuentes-Flores A, Geronimo-Olvera C, Girardi K, et al. Senescent Schwann cells induced by aging and chronic denervation impair axonal regeneration following peripheral nerve injury. EMBO Mol Med. 2023;15(12). doi:10.15252/emmm.202317907

32. Ronchi G, Cillino M, Gambarotta G, et al. Irreversible changes occurring in long-term denervated Schwann cells affect delayed nerve repair. J Neurosurg. 2017;127(4):843–856. doi:10.3171/2016.9.JNS16140

33. Agoston D V, Vink R, Helmy A, Risling M, Nelson D, Prins M. How to Translate Time: The Temporal Aspects of Rodent and Human Pathobiological Processes in Traumatic Brain Injury. J Neurotrauma. 2019;36(11):1724–1737. doi:10.1089/neu.2018.6261

34. Wilcox MB, Laranjeira SG, Eriksson TM, et al. Characterising cellular and molecular features of human peripheral nerve degeneration. Acta Neuropathol Commun. 2020;8(1):1–17. doi:10.1186/s40478-020-00921-w

