## Supplementary material for "Inferior Reinnervation of Reverse End-to-Side Nerve Transfer in a Delayed Nerve Repair Rat Model": Table 1

**Effects of acute vs delayed RETS nerve transfer**

|  | **Acute** | **Delayed** |
| --- | --- | --- |
| Onset latency (FN to RF) | 3.52 ± 0.39 ms | 1.11 ± 0.08 ms |
| Peak amplitude (FN to RF) | 0.75 ± 0.21 mV | 4.28 ± 0.4 mV |
| Onset latency (ON to RF) | 2.50 ± 0.18 ms | 2.74 ± 0.37 ms |
| Peak amplitude (ON to RF) | 1.04 ± 0.19 mV | 0.41 ± 0.07 mV |
| Retrogradely labeled motor neurons (Micro-emerald) | 23 ± 12 (6.6%) | 143 ± 10 (41.1%) |
| Retrogradely labeled motor neurons (Micro-ruby) | 53 ± 8 (54.1%) | 19 ± 4 (19.4%) |

**: p<0.01; ***, p<0.001; ****, p<0.0001, One-way ANOVA as shown in graph

**Number of denervated and reinnervated NMJs**

|  | **Acute** | **Delayed** |
| --- | --- | --- |
| Denervated | 2 ± 1 | 10 ± 2 |
| Innervated by GFP axons | 159 ± 38 | 14 ± 5 |
| Innervated by NF axons | 585 ± 70 | 457 ± 42 |

*: p<0.05, two-way ANOVA as shown in group

**Effects of nerve babysitting**

|  | **ON cut** | **ON cut + PW** | **Delayed RETS** |
| --- | --- | --- | --- |
| Onset latency (FN to RF) | 0.87 ± 0.1 ms | 1.15 ± 0.08 ms | 1.06 ± 0.08 ms |
| Peak amplitude (FN to RF) | 5.52 ± 0.82 mV | 5.53 ± 0.95 mV | 6.31 ± 0.84 mV |
| Retrogradely labeled motor neurons (Micro-ruby) | 173 ± 13 (49.7%) | 150 ± 15 (43.1%) | 151 ± 10 (43.4%) |

**Effect of acute repair vs delayed repair in complete denervated nerve**

|  | **Acute** | **Delayed** |
| --- | --- | --- |
| Onset latency (ON to RF) | 1.77 ± 0.28 ms | 2.17 ± 0.06 ms |
| Peak amplitude (ON to RF) | 1.53 ± 0.40 mV | 0.92 ± 0.19 mV |
| Retrogradely labeled motor neurons (Micro-ruby) | 77 ± 14 (78.5%) | 61 ± 11(62.2%) |

**Table 1**

Quantitative data showing onset latency, peak amplitude and motor neuron count in all groups. Numbers inside parentheses for motor neuron count are percentage of counted cells in total number of motor neuron pool at the level of labeling. FN, femoral nerve; ON, obturator nerve; RF, rectus femoris; PW, perineurial window.
