## Supplementary material for "Inferior Reinnervation of Reverse End-to-Side Nerve Transfer in a Delayed Nerve Repair Rat Model": Supp Fig 1

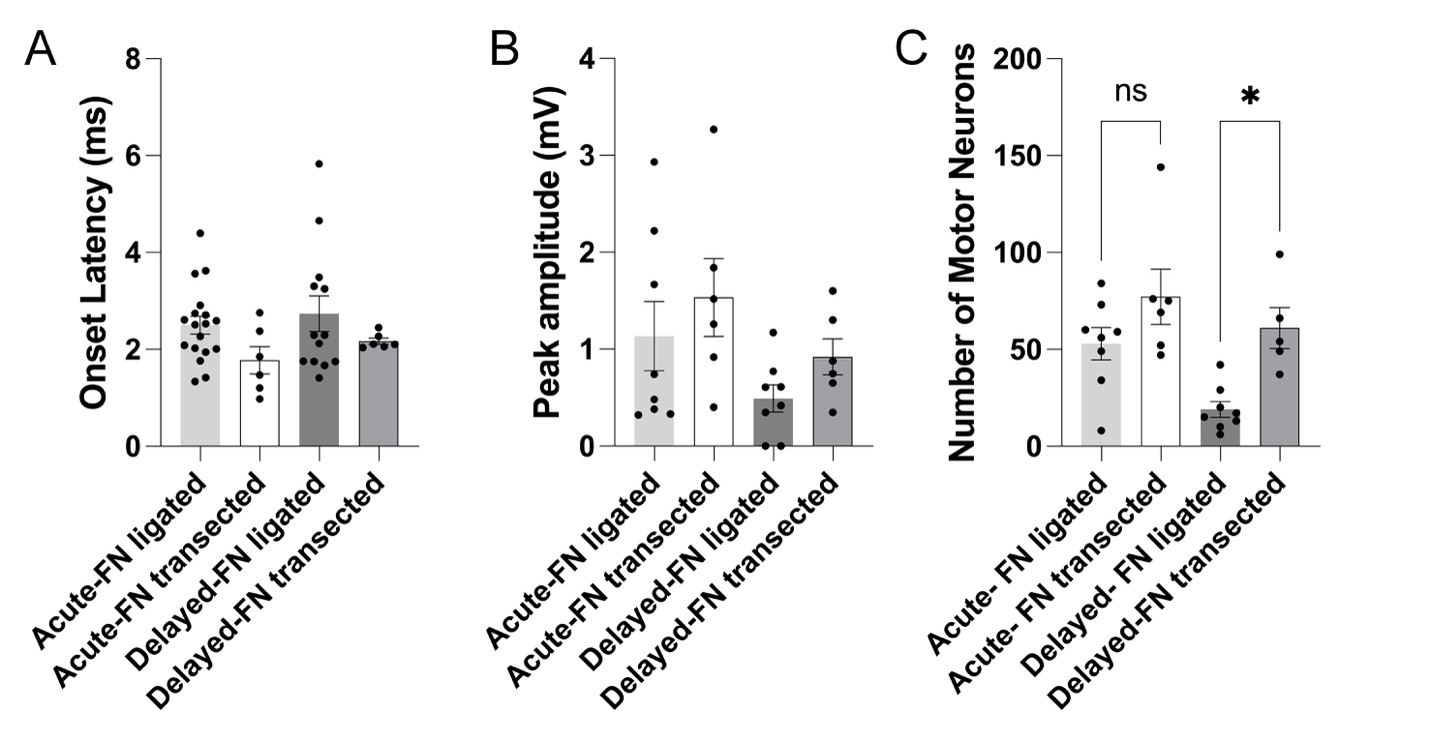

**Supp Fig 1**

Post-hoc analysis of FN crush and double ligation with FN transection showed a trend of shorter latency (A) and higher amplitude (B) but significantly higher number of donor nerve pool motor neurons regenerated into recipient FN in complete (transected) denervated FN compared to partial (double ligated) denervation (C).
